# Fluorescence Lifetime Unmixing: A New Workflow for FLIM Live-Cell Imaging

**DOI:** 10.1101/2025.03.05.641717

**Authors:** Cornelia Wetzker, Marcelo Leomil Zoccoler, Svetlana Iarovenko, Chukwuebuka William Okafornta, Anja Nobst, Hella Hartmann, Thomas Müller-Reichert, Robert Haase, Gunar Fabig

## Abstract

Fluorescence lifetime imaging microscopy (FLIM) translates the duration of excited states of fluorophores into lifetime information as additional source of contrast in images of biological samples. This offers the possibility to separate fluorophores particularly beneficial in case of similar excitation spectra. Here, we demonstrate the distinction of fluorescent molecules based on FLIM phasor analysis, called lifetime unmixing, in live-cell imaging using open-source software for analysis. We showcase two applications using *Caenorhabditis elegans* as a model system. First, we unmixed the highly spectrally overlapping fluorophores mCherry and mKate2 to distinctively track tagged proteins in six-dimensional datasets to investigate cell division in the developing early embryo. Second, we unmixed fluorescence of tagged proteins of interest from masking natural autofluorescence in adult hermaphrodites. For FLIM data handling and workflow implementation, we developed the open-source plugin napari-FLIM-phasor-plotter to implement conversion, visualization, analysis and reuse of FLIM data of different formats. Our work thus advances technical applications and bioimage data management and analysis in FLIM microscopy for life science research.

## INTRODUCTION

Fluorescence microscopy is broadly applied for the investigation of molecular processes in living samples. Largely based on spectrally distinct endogenous or extrinsic fluorophores, it enables the relatively precise localization of subcellular structures in space and time. Due to the spectral overlap of excitation and emission spectra of fluorophores and the limitation of available laser lines and tunable laser ranges, the number of simultaneously investigated cellular features is technically restricted. However, multiplexed simultaneous imaging of multiple cellular features is crucial for an understanding of the complex interplay of biomolecules in cells and tissues.

These limitations can be overcome partially by the application of specialized imaging techniques such as spectral unmixing. Based on dispersion of emitted light, the latter can be used in cases of minimal differences in either excitation or emission spectra or both^1^. However, the technique requires adequate single fluorophore controls. The assessment of fluorescence lifetime in bioimages, termed fluorescence lifetime imaging microscopy (FLIM), is an alternative approach. In the past decades, FLIM has been mostly applied to investigate physical environments of fluorophores, aspects of cellular metabolism and also protein-protein interactions across various species and biological samples^2,3,4^. It gains further importance for the separation of different fluorophore signals in fixed samples which we term lifetime unmixing^5,6,7^.

The fluorescence lifetime being the duration of the excited state upon photon absorption by the fluorochrome is a characteristic attribute of each fluorophore in its microenvironment^2^. This source of contrast can be used to classify pixels or objects in bioimaging. A disadvantage of FLIM is the requirement for higher photon counts for an accurate lifetime decay curve representation and analysis which may require extended image acquisition partially at the cost of temporal and spatial resolution^8,9^. Yet, advancements in laser technology, steady improvement in the sensitivity of detectors and an increase in the number of pulsed lasers in light microscopy facilities have contributed to a higher popularity of this technique and a demand for improved data handling and analysis workflows suitable for specific applications.

Phasor analysis represents an alternative mode of lifetime decay analysis to the broadly applied exponential fitting^10,11,12^. It generates graphical representations of lifetime characteristics per pixel of an image by Fourier transformation of decay data into two-dimensional (2D) plots without the investigator’s assumption of the number of lifetime components of the sample in the experimental scenario. Data points from pixels of similar biological content form clusters in phasor plot visualizations and allow classification of corresponding pixels as one entity for further analysis. In addition, phasor analysis allows the estimation of absolute lifetimes or relative changes between clusters on calibrated setups as well as relative contributions to clusters. Furthermore, phasor analysis is convenient to assess the complexity of distinct fluorophores and fluorochrome states within a given sample^13,5,14,6^. Those diverse workflows make FLIM combined with phasor analysis a powerful imaging technology to explore both known and unknown biomolecules in more depth.

In this work, we demonstrate the use of lifetime unmixing to separate signals of fluorophores of nearly identical emission spectra for imaging of dynamic processes involving conversion and processing of multi-dimensional FLIM datasets. We investigated both early embryos and adult hermaphrodites of the nematode *Caenorhabditis elegans* to showcase two applications of multi-channel phasor FLIM.

To date, the availability of and access to workflows for handling, analysis and sharing of FLIM image data is limited and partially restricted by proprietary file formats^15^. This limits the broad exploration, critical evaluation and reuse of valuable scientific content for the life scientist community. We thus prioritized the use of open-source software which resulted in the development of a plugin for the 3D visualization tool napari, termed napari-flim-phasor-plotter (NFPP, github.com/zoccoler/napari-flim-phasor-plotter/) to visualize and phasor analyze FLIM datasets of various formats^16,17^.

Based on the resulting specific lifetime information clusters of phasor plots, image datasets were processed and analyzed using the broad napari plugin ecosystem along with standard open-source image processing libraries. The plugin further includes the conversion of FLIM data to the chunked file format Zarr for handling N-dimensional data arrays^18^. The Zarr format allows lazy loading for faster visualization and processing using cloud-based tools. Moreover, we present a practicable workflow to visualize, share and publish 6D FLIM datasets in the bioimage management platform Open Microscopy Environment (OME) Remote Objects (OMERO)^19,20^.

## MATERIALS AND METHODS

### *C. elegans* maintenance

*C. elegans* strains (Table 1) were grown on nematode growth medium (NGM) plates at 20°C with *E. coli* (OP50) as food source according to established protocols^21^.

**Table 1:**
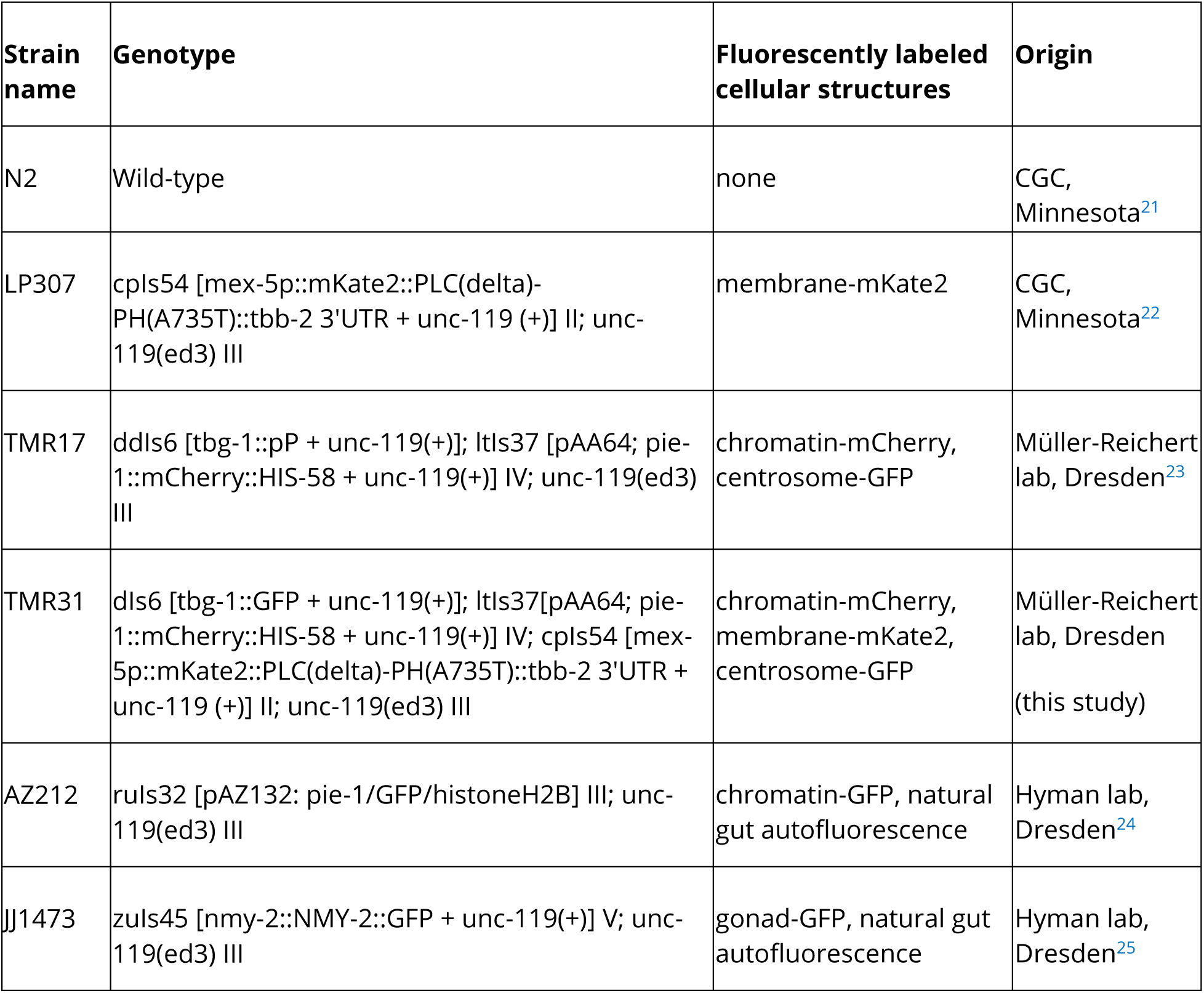
Transgenic *C. elegans* lines.

For imaging of *C. elegans* embryos, lines single- or double-positive for fluorescently tagged proteins, namely an mCherry-tagged chromosome marker (chromatin-mCherry, HIS-58), an mKate2-tagged membrane marker (membrane-mKate2, phospholipase C delta) and a GFP-tagged centrosome marker (centrosome-GFP, γ-tubulin chain 1), and a wild-type negative line (N2) were used^26,22,27^. For imaging of hermaphrodites, lines expressing a GFP-tagged chromosome marker (chromosome-GFP, histone H2B) or a GFP-tagged gonadal marker (gonad-GFP, NMY-2) were used.

### Sample preparation for imaging

For imaging of early *C. elegans* embryos, adult hermaphrodites were dissected in M9 buffer containing 25 mM levamisole hydrochloride (MP Biomedicals, Solon, USA). Isolated embryos were mounted on a 4% (w/v) agarose pad between a slide and a coverslip^28,29^. For visualization of whole animals, we immobilized the worms in between an agar pad and a coverslip using M9 buffer containing 25 mM levamisole hydrochloride^30,31^.

### FLIM microscopy

FLIM was performed on the basis of time-correlated single photon counting (TCSPC) using an upright confocal laser scanning microscope (STELLARIS 8, Leica Microsystems, Germany) with a pulsed supercontinuum laser (NKT Photonics, Germany) for single photon excitation at 40 MHz frequency using a water immersion objective (40x/1.1, HC PL APO CS2, Leica Microsystems, Germany)^32^. We used the acquisition software Leica Application Suite X (LAS X) (version 4.5.0.25531) including the FLIM/FCS license (version 4.5.0). GFP was excited at 480 nm (approximately 16 µW in none-objective control) and emitted in the range of 500 to 560 nm. mCherry and mKate2 were excited either alone or simultaneously at 590 nm (approximately 84 µW in none-objective control) and emitted in the range of 610 to 800 nm. The pinhole was set to one Airy unit at 600 nm reference. Both spectral ranges were imaged simultaneously. Emitted light was detected with hybrid HyD X detectors (Leica Microsystems, Germany).

The field of view and zoom factor of 4 were chosen to image whole embryos as z stacks of FLIM images of 256 x 256 pixels. The image stacks were recorded with 600 Hz speed of bi-directional scans and optical zoom of 4. This yielded a pixel dwell time of 2.4 µs, a frame time of 0.66 s and voxel sizes of 0.27 µm in x and y and 0.5 µm in the z dimension. To collect sufficient photons for FLIM imaging, three frames were summed per image yielding a total scan time of approximately 40 s or shorter per image stack. Acquisition was performed continuously from two-cell stage prior to the first division until condensed chromatin in the second division appeared (differing from 18 to 30 min between embryos).

For whole worms we recorded several time points but analyzed only a single. Two channels z stacks of FLIM images were acquired with the same setup as embryos. Differences in central settings included optical zoom of 1, image sizes of 512 x 128 pixels resulting in voxel sizes of 0.54 µm x 0.54 µm x 0.5 µm in x, y,z. Pixel dwell times were 1.2 µs and frame times of 0.58 s with two added frames leading to 99.2 s (AZ212) and 116.5 s (JJ1473) per stack per time point. The 6D FLIM image datasets were then saved in LIF format in a containerized data structure. We then exported 6D embryo datasets and 5D single time point adult datasets as individual 4D files in PTU format (dimensions being x, y, channels and photon counts in the FLIM dimension which we refer to as microtime) for each z plane and time point as sole export mode for downstream data processing and sharing.

### Scans for excitation spectra of fluorophores

Spectral excitation scan images of one biological replicate per genotype were acquired on the same microscope with same setup as FLIM images in the range of 440 to 790 nm with 67 steps of 5 nm step size using the white-light laser (wavelength-dependent power between 13 and 51 µW in none-objective control) at 40 MHz pulse rate. Emitted photons were detected in a 20 nm spectral window of 10 nm distance of excitation wavelength. Datasets were saved along FLIM images in the corresponding LIF files prior to export. Photon counts were summed and normalized across whole images for each series and plotted for comparison of excitation spectra of fluorophores.

### Conversion and storage of FLIM image formats

We developed the napari FLIM phasor plotter (NFPP) plugin for the open-source image viewer napari to re-build each 6D dataset as a single stack from multiple individual 4D PTU files and convert them to single Zarr (https://zarr.dev/) arrays. Each file contained the following dimensions: channel, photon counts per time bin (microtime), time, z, y and x. This conversion to Zarr divides N-dimensional datasets into so-called chunks. This allowed lazy loading, an optimized loading into memory of the subset of data needed at a time for increased speed and working with larger-than-memory data sizes, into napari. Conversion to Zarr further improved visualization and downstream processing of datasets, particularly beneficial for large datasets.

We deposited these imaging datasets in an instance of the bioimage management software OMERO to simplify annotation, collaboration between internal project partners and sharing of data with the research community. The generated datasets and associated metadata are accessible on a public user space of the OMERO instance (https://omero.med.tu-dresden.de/webclient/?show=project-1401)^33^. Raw and processed bioimage datasets will further be made available for reuse in a public biological image repository.

Currently, OMERO upload and visualization are supported for datasets up to 5 dimensions (5D). Hence, visualization of 6D datasets using open-source tools requires several steps of reformatting, including reduction to 5D for sharing via OMERO. We transformed 6D Zarr to 5D OME-TIFF datasets by conversion including separation of individual time points. Here, the time dimension of files stores the lifetime decay information. This allowed OMERO upload and thus visual exploration of the lifetime dimension with a slider via the OMERO web interface.

Finally, we also built an extra single OME-TIFF file per dataset with the microtime dimension reduced to summed intensity information and time points of the actual time series in the time dimension. Hence, these 5D intensity datasets lacked the fluorescence lifetime information but allowed continuous navigation throughout the whole time-lapse using a slider with the option of adding segmentation results as an extra channel in OMERO (see example dataset, accessed via https://omero.med.tu-dresden.de/iviewer/?images=37177&dataset=2037). Bioimage datasets were annotated according to REMBI recommendations^34^.

### Lifetime unmixing based on FLIM phasors

We distinguished the fluorescence signal of the chromatin label mCherry based on its lifetime signature distinct from that of mKate2 using NFPP. The plugin uses the napari-clusters-plotter (https://github.com/BiAPoL/napari-clusters-plotter) for phasor visualization. We defined a region in the phasor plot that corresponded exclusively to mCherry labeled voxels. This region was selected manually based on the strain TMR17, which contained only mCherry-labeled histone H2B. We defined the phasor lifetime signature of mCherry-tagged proteins by selection of an elliptic field around the single cluster that was broad enough to always encompass centers of clusters over all time-points but not too large to decrease noise inclusion. This fixed elliptic region was then applied to phasor plots of FLIM datasets from *C. elegans* strains with different tagged proteins in the red spectrum (double positive for mCherry and mKate2, and single positive for mKate2).

This workflow is available in the GitHub repository (https://github.com/zoccoler/Lifetime-Unmixing/, in the python script ‘phasor_plot_analysis_batch_processing.py’). Briefly, each dataset converted to the Zarr format, available as an option for large datasets in NFPP, was loaded into napari. Here, phasor plots were generated with a fixed set of parameters and the fixed, manually adjusted ellipse selection was applied to generate a new layer of segmentation mask resulting from the selection. To reduce noise, morphological opening, fill holes and minimal object size filtering methods were applied on the masks^35,36,37^. Finally, instance segmentation retrieved individual objects representing the chromatin, followed by feature extraction to get each object’s centroid position and volume for downstream analysis of chromatin dynamics.

This segmentation workflow generated the following output: a new channel complementary to the intensity image over the whole time-lapse as OME-TIFF files, phasor and pixel coordinates tables for each time-point as CSV files, phasor plot screenshots before and after selection for each time-point as PNG files, objects centroid and volume tables over the whole time lapse as CSV files, and parameters values used as TXT files. These results are available via OMERO. The position of the centroids of the segmented chromosomes was used to calculate the three-dimensional (3D) Euklidean distance among them in each frame. Then the distances were plotted over time and an exponential function was fitted to the measurements using Python to estimate the initial speed of chromosome segregation^38^.

For the analysis of the pole-to-pole distance, the centrosomes were segmented using the GFP fluorescence channel by adaptive local thresholding^39^. Then the distance was calculated analogously as described above. A polynomial 1D function was fitted to the pole-to-pole distance from anaphase onset (0 s) to 120 s after the initiation of anaphase as this period resembled the linear part of the sigmoid curve. From this, the speed of pole-to-pole separation could be estimated.

### Analysis of adult worms

We analyzed worm lines with GFP-tagged proteins, namely histone H2B (strain AZ212) and non-muscle myosin-2 (NMY-2, strain JJ1473). The histone H2B-tagged strain AZ212 is used to follow chromatin dynamics in both gonadal and somatic cells in *C. elegans*. Non-muscle myosin-2 is expressed in the worm gonad and important in cell division^40^. Present in the gonad of hermaphrodites, it is involved in maintaining the gonads’ structure and the oocyte formation^41^.

A common problem when imaging living animals such as *C. elegans* is autofluorescence, mostly associated with the worm’s intestine^42,43^. In these examples, the lifetime of pixels representing autofluorescence and GFP are quite different, which implies clearly distinct clusters in the phasor plots.

We generated phasor plots of FLIM datasets and automatically segmented data points of the phasor plot into two distinct clusters using the K-Means clustering algorithm from the scikit-learn python library through the NFPP plugin^44,45^. We saved the voxels of each cluster as a separate pseudo-channel and uploaded these results to OMERO. For identification of cell nuclei we used two segmentation workflows: (a) we applied the Voronoi-Otsu labeling algorithm from pyclesperanto (https://zenodo.org/records/10432619), counted the number of segmented objects, and stored their centroid positions and volume in a table; (b) we used fill holes and morphological operations (area_closing, isotropic_closing and isotropic opening) on the pseudo-channel representing the mCherry histone H2B cluster^46^. Then we applied again a Voronoi-Otsu labeling, counted the number of objects, and stored their centroid positions and volumes as well. Both workflows can be visualized and reproduced by running the following notebook in the code repository: https://github.com/zoccoler/Lifetime-Unmixing/blob/main/code/phasor_plot_AZ212.ipynb. We uploaded the segmentation results as new channels to OMERO along with the number of objects to compare the improvement of accuracy after lifetime unmixing.

### Data presentation

For the generation of figures (Figs. 1–4), we used OMERO.figure (https://www.openmicroscopy.org/omero/figure/). This provides access to the original and processed datasets directly through hyperlinks in the composed figures. For depiction according to conventions, images of embryos were rotated in a stereotypical way with the larger (AB) cell always positioned on the left and the smaller (P1) cell always on the right.

**Fig. 1:**
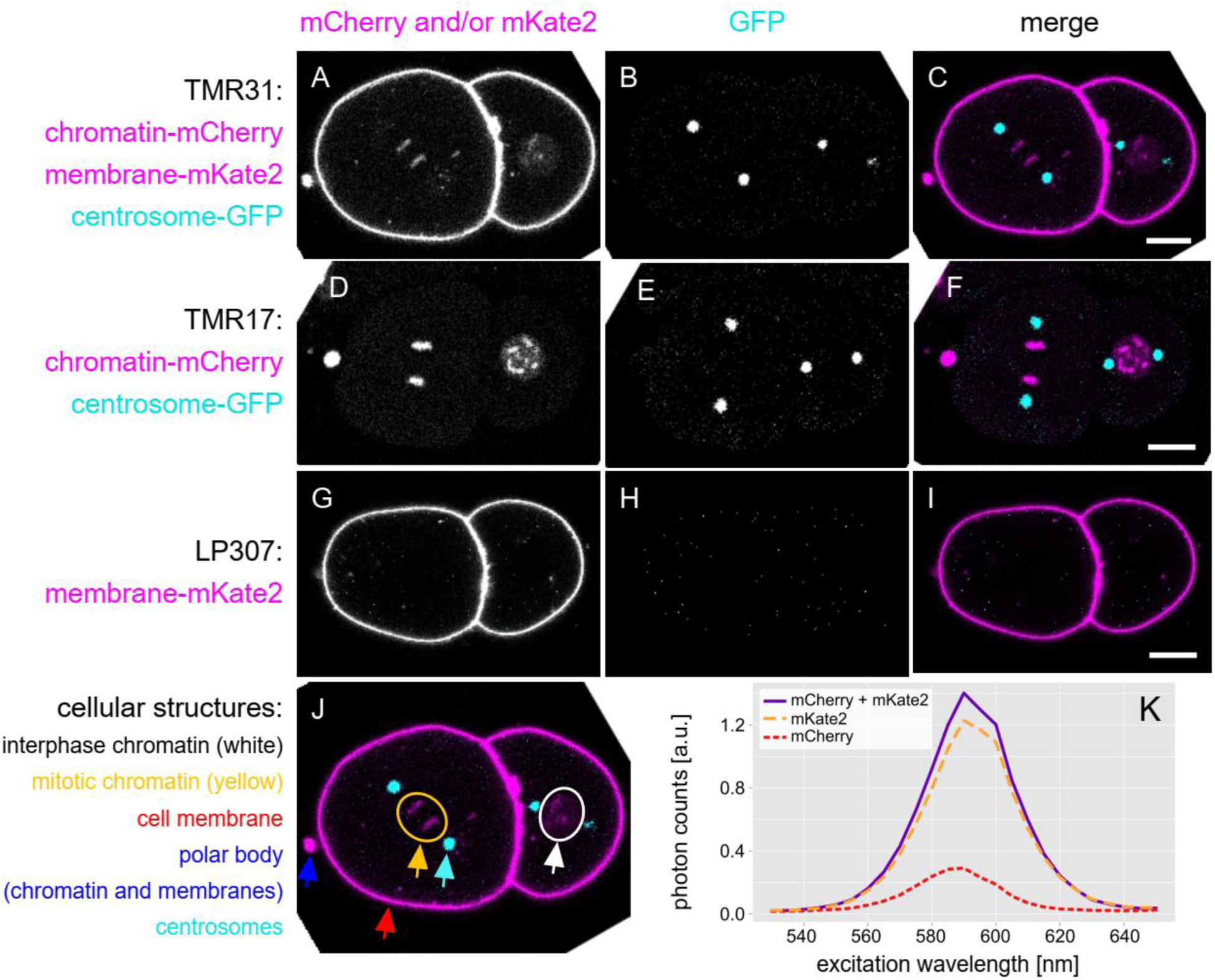
Intensity imaging of worm strains and excitation scans demonstrate the spectral similarity of the red fluorophores mKate2 and mCherry and cellular localization in embryos. (**A-J**) Single images of summed intensity stacks of *C. elegans* lines single- or double-positive for chromatin-mCherry and membrane-mKate2 show localization of fluorescence to chromatin and membranes, respectively, in 2-cell embryos. Centrosome-GFP further marks the mitotic spindles (columns 2 and 3). (**J**) Triple fluorescent embryos highlight interphase (white ellipse) and mitotic chromatin (yellow ellipse), cell membranes (red arrow), a polar body (blue arrow) with dense co-presence of chromatin and membrane and centrosomes (cyan arrow). Scale bars, 10 µm. (**K**) Excitation scans in the range of 530 to 650 nm in single- and double-positive embryos demonstrate spectral overlap of fluorescence of mCherry and mKate2. (Link to Figure 1: https://omero.med.tu-dresden.de/figure/file/5748/)

## RESULTS

### Characterization of fluorophores with overlapping spectra

To visualize dynamics of cellular processes during cell divisions,we imaged *C. elegans* embryos simultaneously expressing chromatin-mCherry, membrane-mKate2 and centrosome-GFP. Asboth fluorophores, mCherry and mKate2 emit in the red light spectrum, there was substantial spectral overlap limiting the number of simultaneously detectable macromolecules. We first characterized the spectral overlap of red fluorophores by image visualization in conventional confocal microscopy mode as 3D intensity image stacks at a single time point (Fig. 1, https://omero.med.tu-dresden.de/figure/file/5748/). The mCherry-tagged protein labeled indeed the chromatin of cells in either interphase or mitosis (Fig. 1 A-F, J), while the mKate2-tagged protein marked the membrane of the embryonic cells (Fig. 1 A-C, G-I, J). Both histone-mCherry and membrane-mKate2 fluorescence were detected also at the extruded polar bodies (Fig. 1 J) In addition, we confirmed the labeling of the spindle poles by centrosome-GFP (Fig. 1 A-F, J). Fluorescence imaging further revealed a difference in local intensity of the chromatin-mCherry *versus* the membrane-mKate2 signal. The chromatin signal was much weaker compared to the membrane signal, an observation we could confirm by spectral scans (Fig. 1 K). Emission scans across a broad excitation range confirmed the spectral similarity of mCherry and mKate2 fluorescence in the imaged worms.

### Lifetime unmixing of spectrally similar fluorophores using the napari-FLIM-phasor-plotter (NFPP) and napari

To demonstrate the applicability of the NFPP plugin in live-cell imaging, we performed lifetime unmixing of mCherry and mKate2-tagged proteins during mitotis in early *C. elegans* embryos. Using NFPP, we separated the superimposed red fluorescence signals of the chromosomes and of the membranes into fluorophore-specific pseudo-channels. Here, NFPP was used for the import, visualization and processing of multi-dimensional FLIM datasets of several conventional formats including SDT, PTU and TIFF. The plugin thus extends the N-dimensional python-based image viewer napari, a powerful community-driven open-source platform for image visualization and analysis, to FLIM datasets.

To separate FLIM datasets into distinct pseudo-channels, we generated 6D FLIM datasets of xyzctµ dimensions, being x, y and z spatial dimensions, as well as dual channels (c), a time-series (t) and FLIM microtime (µ) dimension (Fig. 2, https://omero.med.tu-dresden.de/figure/file/5749/). We show intensity images of maximum projections of single time-point sub-stacks of early developing *C. elegans* embryos carrying single or double red fluorophores (Fig. 2 A-C).

**Fig. 2:**
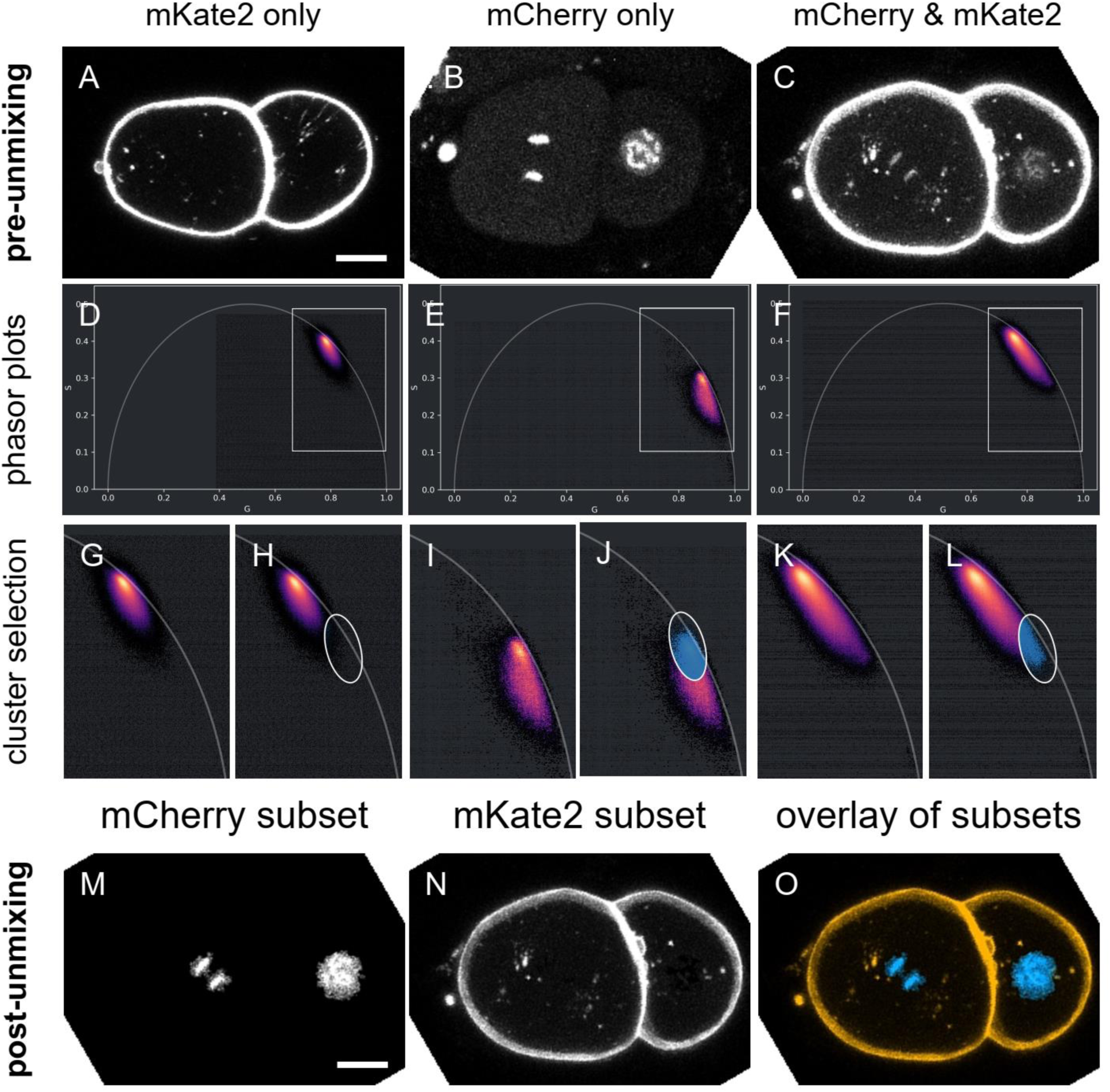
Separation of image stacks into distinct sub-datasets of spectrally similar fluorophores by lifetime unmixing. (**A**, **B**) Sub-stack of images of an embryo tagged with mKate2 or mCherry only and (**C**) an embryo with mCherry and mKate2 signals. (**D** - **F**) Corresponding full lifetime phasor plots. (**G - L**) Regions of interests of the respective phasor plots of (**D - F**) with with ellipse drawn to select a subset of pixels for lifetime unmixing (**H, J, L**). The double-labeled images (**C**) were further processed. (**M**) Lifetime unmixed data set highlighting the mCherry-tagged chromatin. (**N**) Lifetime unmixed data set highlighting the mKate2-tagged cell membrane. (**O**) Lifetime unmixed data set corresponding to (**C**) with overlaid chromatin (blue) and cell membrane (magenta). The unmixing was based on the selected signals from the phasor plot (**L**). Scale bar, 10 µm. Sub-stack planes 19-28, time-point 6 (**A**), planes 16-25, time-point 15 (**B**) and planes 19-28, timepoint 10 (**C**). (Link to Figure 2: https://omero.med.tu-dresden.de/figure/file/5749/)

The exported data of the acquisition system was then imported into napari using NFPP. The plugin further converted the lifetime information per voxel into a 2D plot termed phasor plot by Fourier transformation (Fig. 2 D-F). Magnifications of regions of interest of phasor plots emphasize the difference in lifetime-based localization of fluorophore-specific data points visualized as density plots (Fig. 2 G, I, K) that are a prerequisite for successful lifetime unmixing of datasets by selection of phasor clusters (Fig. 2 H, J, L). Due to the high abundance of mKate2-positive voxels, the signal of mCherry-positive data points is barely visible in the phasor plots (Fig. 2 F, K, L) which emphasizes the importance of single fluorophore controls (Fig. 2 D, G, H and Fig. 2 E, I, J). Overall, red fluorescence of distinct fluorophores could be separated into sub-datasets of chromatin and membrane signal by lifetime unmixing using NFPP (Fig. 2 M-O).

### Lifetime unmixing for live imaging of cell division

Next, we investigated the dynamics of chromosomes and centrosomes during mitosis in two-cell *C. elegans* embryos by downstream analysis of lifetime-unmixed datasets in napari (Fig.3, https://omero.med.tu-dresden.de/figure/file/5819/). For that the aforementioned method was used to process the raw data (Fig. 3A) and segment the mCherry-positive voxels in the phasor plot (Fig. 3B). Data was then filtered, resulting in much less segmented voxels (Fig. 3C) which enabled a selection of those segments containing the chromosomes (Fig. 3D). We then calculated the centers of mass of each segment (Fig. 3D, white crosses) and used these coordinates to determine the chromosome-to-chromosome distance over time (Fig. 3E). By fitting an exponential curve we determined a speed of 11.64 µm/min for the chromosomes in the larger AB cell and a speed of 13.21 µm/min in the smaller P1 cell of the two-cell stage embryo. For centrosomes, we used adaptive local thresholding for segmentation and used the resulting center coordinates to calculate the pole-to-pole distances over time (Fig. 3F). By fitting a linear function to the pole-to-pole distance after anaphase onset until 120 s after anaphase onset, we determined a speed of pole separation of 3.36 µm/min for the AB cell and 2.78 µm/min for the P1 cell. These values matched previously published data on centrosome separation in the two-cell embryo^46^. Hence, we demonstrated that lifetime unmixed live-cell bioimages can be applied to precisely track dynamic processes during animal development.

**Fig. 3:**
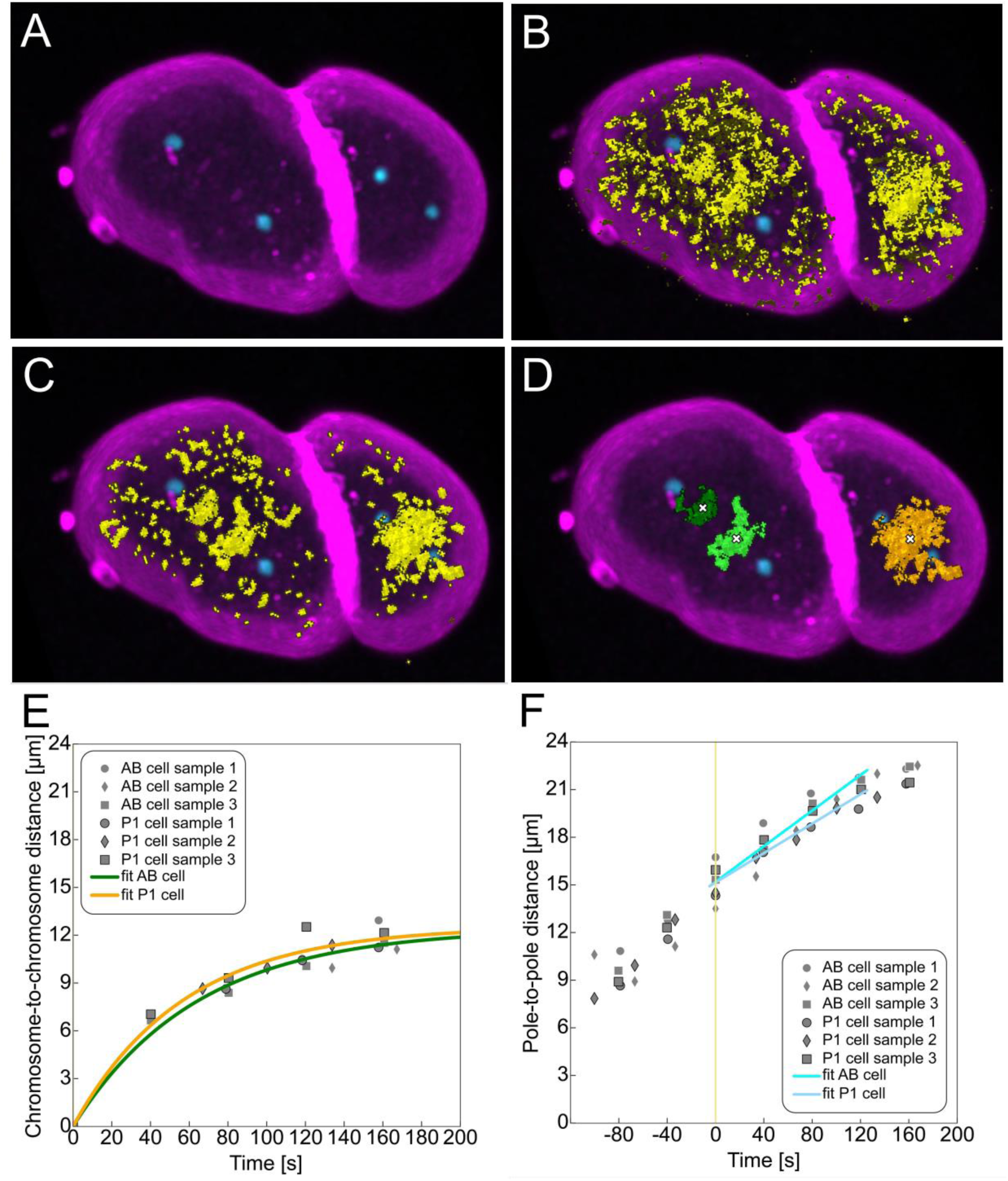
Workflow from lifetime unmixed mCherry channel using the NFPP plugin for analysis of spindle dynamics. (**A**) Maximum intensity projection of a confocal image stack of one TMR31 *C. elegans* embryo (timepoint 10) double positive for chromatin-mCherry and membrane-mKate2 (both in magenta) combined with centrosome-GFP (cyan), 3D rendered with napari. (**B**) Same image as in (A) overlayed by a yellow mask resulting from ellipse cluster selection from the phasor plot. (**C**) Image with phasor mask post-processed by the NFPP plugin function “smooth cluster mask”: fill holes (scikit-image “area closing”) + connect nearby objects (scikit-image “isotropic closing”) + remove small objects (scikit-image “isotropic opening”). (**D**) Instance segmentation on post-processed phasor mask from (C) by connected components labeling (scikit-image “label”) followed by removing objects smaller than volume threshold (pyclesperanto prototype “exclude small labels”). The white crosses indicate the center of the segments used for the determination of the distance of chromosomes over time. For scale, the length of the *C. elegans* embryo along the long embryo axis is about 50 µm. (**E**). Plot showing chromosome-to-chromossome distance over time for the AB- and P1-cell in each analyzed TMR31 embryo (grey symbols) along with exponential curves fitted to the measurements (green and yellow). (**F**) Plot showing pole-to-pole distance over time for the ABand P1-cell in each analyzed TMR31 embryo (grey symbols) along with linear curves fitted to the measurements (cyan and light blue) from anaphase onset (t=0) until 120s after anaphase onset. (Link to Figure 3: https://omero.med.tu-dresden.de/figure/file/5819/)

### Removal of autofluorescence from bioimage datasets by lifetime unmixing

We further evaluated the applicability of the established workflow of lifetime unmixing using the NFPP plugin to split natural autofluorescence from spectrally overlapping GFP fluorescence marking biomolecules of interest in adult worms (Fig. 4, https://omero.med.tu-dresden.de/figure/file/5820/). Strong intrinsic autofluorescence of biological samples is often a challenge in imaging of both tissues and whole organisms and can mask and hinder the analysis of cellular processes of interest. Biomolecules such as chitin, melanin, NADH and FAD and many others, are fluorescent in green or red spectral ranges^48,49,4,50^. Such biomolecules with structural, protective and/or metabolic functions can occur at high concentrations across tissues and also whole organisms^6^. As an example, the gut of worms is highly autofluorescent, thus hindering reliable microscopic analysis of nearby tissues such as the gonad.

**Fig. 4:**
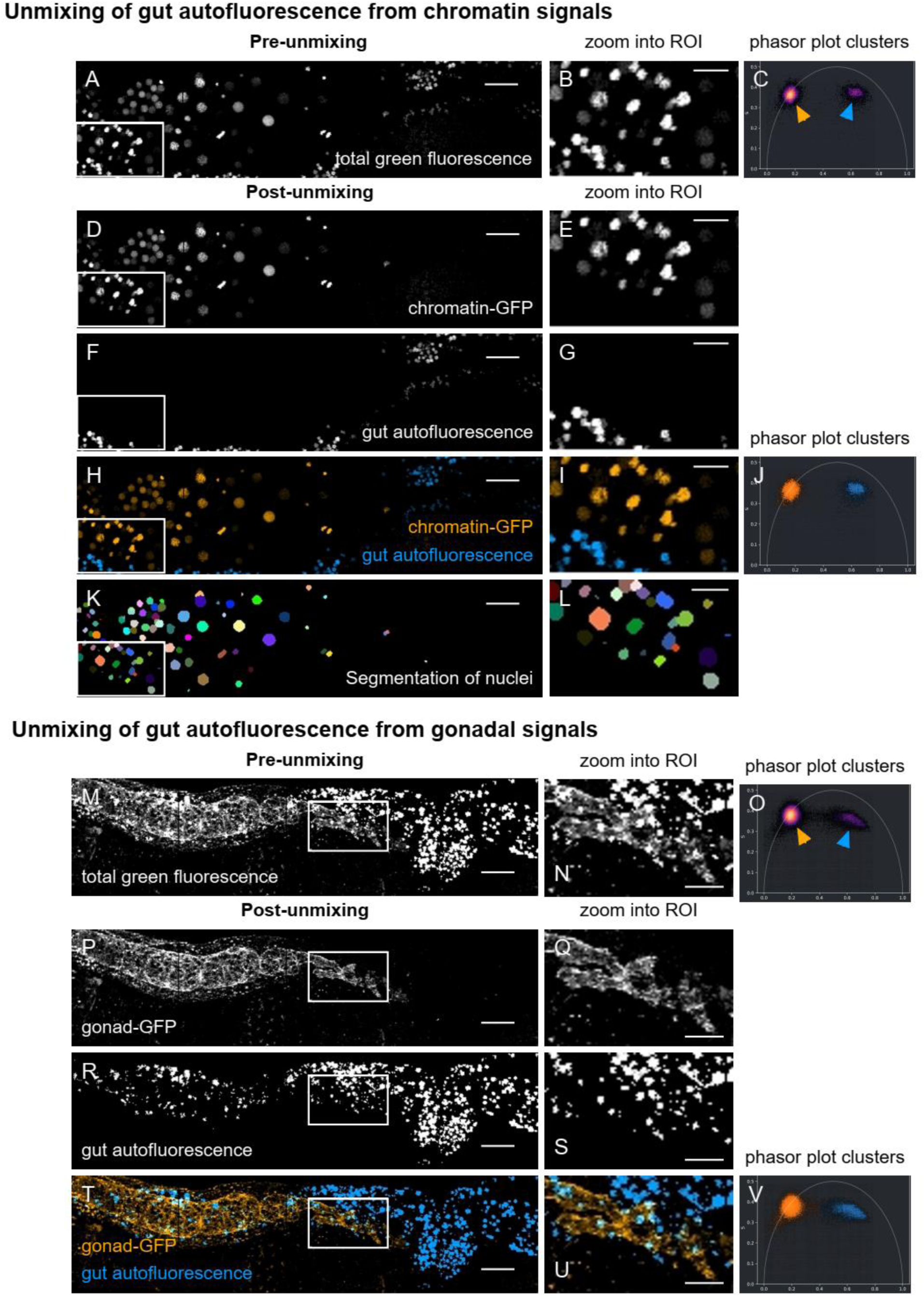
Removal of gut autofluorescence from images of adult C. elegans hermaphrodites by lifetime unmixing. (**A**, **B**) Chromatin-GFP and gut autofluorescence both emit in the GFP spectrum. (**C**) NFPP-created phasor plot showing two distinct populations of fluorescence (orange and blue arrow). Lifetime unmixing creates distinct datasets of GFP-tagged chromatin (orange) and gut autofluorescence (blue) as a basis for pseudo-channel-specific segmentation of cell nuclei. (**D**, **E**) Chromatin-GFP signals separated by lifetime unmixing. (**F**, **G**) Gut autofluorescence separated by lifetime unmixing. (**H**, **I**) Overlay of separated chromatin-GFP signals (orange) and gut autofluorescence (blue). (**J**) NFPP-created phasor plot showing two the two fluorescence populations selected (orange and blue clusters). (**K**, **L**) Segmentation of GFP-positive lifetime unmixed nuclei using napari. (**M**, **N**) Gonad-GFP and gut autofluorescence both emit in the GFP spectrum. (**O**) NFPP-created phasor plot showing two distinct populations of fluorescence (orange and blue arrow). Lifetime unmixing creates distinct datasets of the GFP-tagged gonad (orange) and gut autofluorescence (blue) as a basis for pseudo-channel-specific segmentation of the gonad. (**P**, **Q**) Gonad-GFP signals separated by lifetime unmixing. (**R**, **S**) Gut autofluorescence separated by lifetime unmixing. (**T**, **U**) Overlay of separated gonad-GFP signals (orange) and gut autofluorescence (blue). (**V**) NFPP-created phasor plot showing two the two fluorescence populations selected (orange and blue clusters). Scale bars, 20 µm in overview images (first column) and 10 µm in magnified regions of interest (second column). (Link to Figure 4: https://omero.med.tu-dresden.de/figure/file/5820/)

We imaged adult hermaphrodites with either chromosome-GFP (Fig. 4 A-J) or gonad-GFP (Fig. 4 K-R) fluorescent proteins, both known to be in close proximity to the gut as a source of animal-intrinsic autofluorescence. We could clearly distinguish nuclei of the gonad in the chromosome-GFP worm line. Lifetime unmixing enabled removal of the gut autofluorescence and subsequent segmentation of the gonadal nuclei for potential further analysis (Fig. 4 C, D, I-L). In addition, removal of gut autofluorescence from worms expressing gonad-GFP (Fig. 4 M and N) allowed the distinct localization and segmentation of the membrane and other features of the gonad (Fig. 4 M to R). Compared to conventional intensity imaging, removal of autofluorescence by lifetime unmixing improves the localization of biomolecules of interest and thus the quality of scientific output from live-cell imaging across scales.

## DISCUSSION

FLIM contributes a valuable additional dimension of information to bioimaging beneficial for the identification, characterization and/or separation of fluorophores in their immediate environment. Here, we present lifetime unmixing for the separation of sample-intrinsic and -extrinsic fluorophores in developing live organisms using a newly developed napari plugin, NFPP. This plugin feeds datasets with a lifetime decay dimension into napari, linked to first modes of lifetime analysis and conversion of FLIM datasets to Zarr format for downstream analysis and visualization. Our work emphasizes the current challenges and first solutions for FLIM in general and in 6D datasets in particular. This imaging technique demands a carefully designed acquisition setup combined with enhanced interoperability of data formats and analysis workflows to simplify and standardize its application.

### Acquisition of FLIM data

FLIM requires a careful balance of lifetime, spatial and temporal resolution (Fig. 5). In detail, the detection of sufficient photons is required for precise lifetime decay estimations. Despite the constantly increasing sensitivity of detectors, FLIM requires relatively long image acquisition times compared to corresponding intensity measurements. This challenges particularly multi-dimensional imaging of dynamic processes in living cells as temporal resolution, both in lifetime and time series dimension, and spatial resolution have to be leveled carefully. For example, higher photon counts in the lifetime dimension render decay curves less noisy, leading to more compact and thus distinct phasor clusters and more precise lifetime calculations compared to lower photon count scenarios. Specifically, if the lifetimes of fluorophores are similar, showing differences of less than one nanosecond, unmixing may require a high number of photons. In addition, a phasor cluster of autofluorophores with large variation due to structural variability of the fluorochrome or diverse interaction partners and may thus require a high number of photons.

**Fig. 5:**
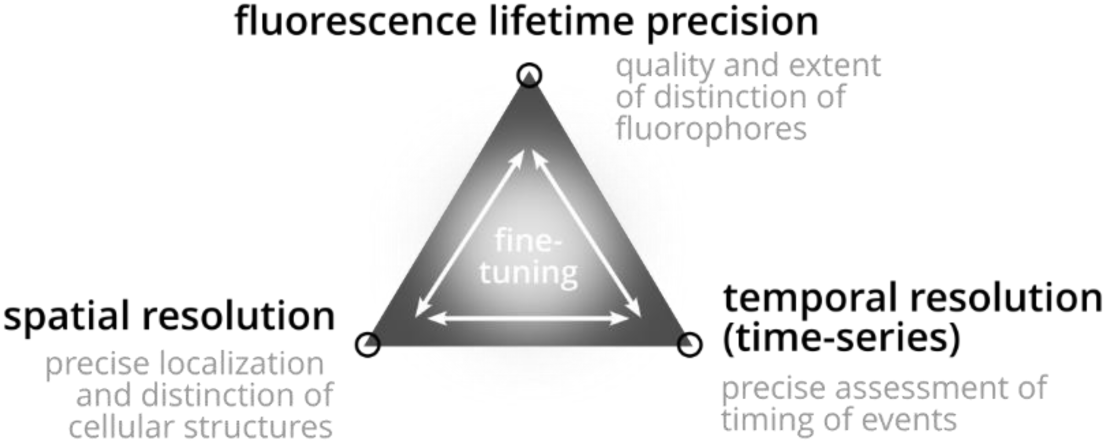
The triangle of compromise in FLIM imaging. Levels of resolution in lifetime, space and time-series resolution require careful and question-specific fine-tuning to obtain the required level of information across all dimensions.

In general, the size of investigated structures defines the required resolution in x, y and z dimensions. Pixel binning might be an option to improve the lifetime precision, if the investigated structures can still be identified. Furthermore, the temporal resolution of a dynamic process is also a critical parameter. As for the investigation of chromatin dynamics in our study, the distinction of individual chromosomes was essential to track their dynamics during separation. However, the density of chromosomal proteins in interphase *versus* mitosis clearly differs, consequently leading to different photon counts across pixels and thus different quality of phasor clusters and pixel classification. In addition, investigation of overall less abundant proteins with spectrally similar tags might further challenge the pixel classification based on phasor clusters. It is, therefore, important to fine-tune the individual steps of the workflow, including data acquisition, lifetime unmixing and image analysis. Only then, FLIM is a powerful imaging technique for the multi-dimensional investigation of dynamic processes.

### Handling and storage of FLIM data

Critical for a broad range of FLIM applications are aspects concerning data handling and analysis as exemplified here for 6D datasets (Fig. 6). First, FLIM data is to a large extent stored in less abundant and thus less supported formats including PTU and SDT. This hinders implementation of proper handling of FLIM data along the data life cycle ranging from import to visualization, analysis, sharing for publication and reuse and archiving. Processing of lifetime information demands specialized workflows, most prominently exponential fitting or phasor analysis based on Fourier transformation of decay information, essential for correlation with further measurements. Thus, FLIM analysis requires extensive integration into image analysis tools with various functionalities to adjust to research questions. First available community-developed open-source tools include FLIMfit (https://github.com/flimfit/FLIMfit/), FlimJ (https://imagej.net/plugins/flimj/) for Fiji, FLUTE (https://github.com/LaboratoryOpticsBiosciences/FLUTE/) and phasor-py (https://www.phasorpy.org/)^51,52^.

**Fig. 6:**
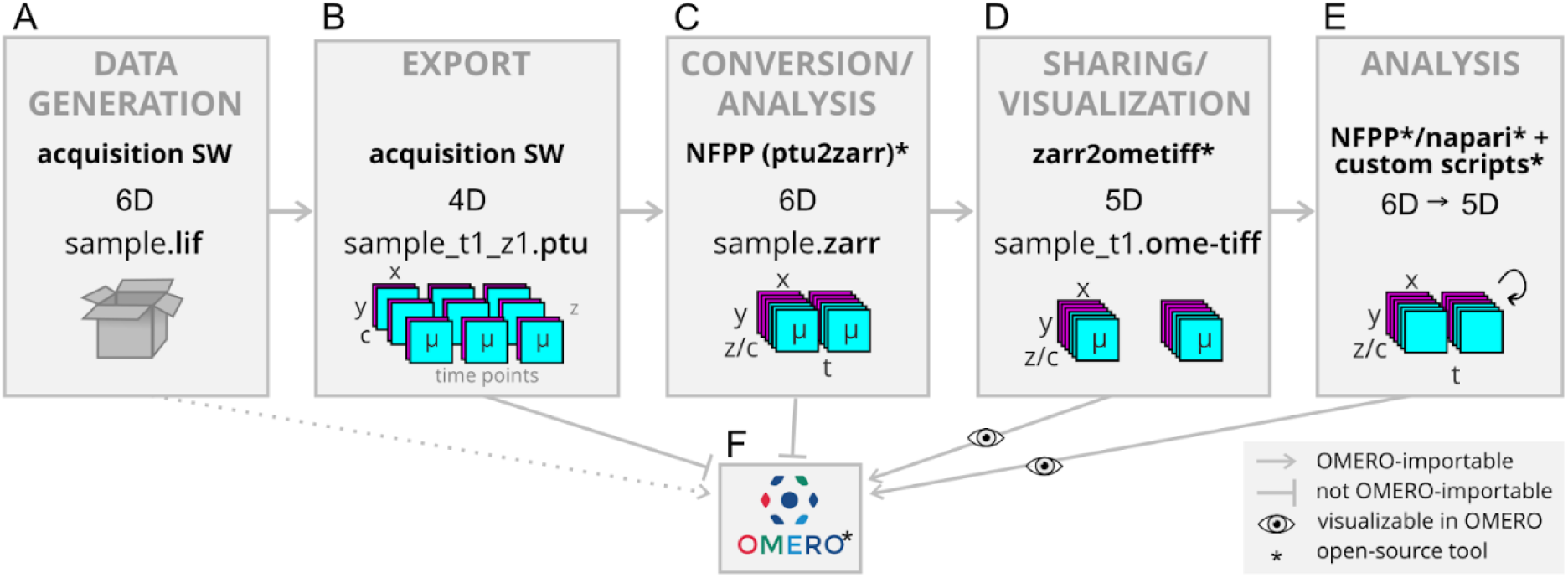
Workflow of FLIM file format conversion for sharing and analysis. (**A**) Storage of 6D FLIM datasets in LIF format using proprietary acquisition software. (**B**) Export of data in 4D sub-datasets in PTU format split by z planes and time points. (**C**) Reassembly of datasets to 6D using the napari-flim-phasor-plotter (NFPP) plugin, and optionally converted to Zarr. (**D**) Conversion to the OME-TIFF format for 5D datasets split by time points permits OMERO upload of image data and metadata. (**E**) Reduction of 6D data in Zarr format to 5D by FLIM analysis using NFPP along with napari and custom scripts contributes additional segmentation masks to 5D datasets in OMERO. (**F**) Overview of possible import of data including segmentation results based on lifetime unmixing into OMERO and possibility of visualization of lifetimes as time series. Lines with arrows indicate the option for OMERO upload of each dataset in its format and number of dimensions. Blocked lines indicate no current option for OMERO upload limited by format (PTU) or 6 dimensions (Zarr). Files in LIF format can be formally uploaded but do not display properly. Eye symbols indicate proper visualization of datasets in OMERO. Asterisks indicate open-source tools and scripts. x and y, x and y spatial dimensions; z, z spatial dimension; c, channel; t, time dimension; µ, photon counts in lifetime dimension (microtime); SW, software; NFPP, napari-flim-phasor-plotter plugin.

In our work, we present a comprehensive approach for 6D FLIM data analysis with increased implementation of FAIR principles to make FLIM datasets more findable, accessible, interoperable and reusable^53^. In our study, datasets were initially stored in LIF format with limited lifetime visualization in OMERO (Fig. 6A, Suppl Fig. 1) and acquisition software-restricted export of only split datasets in PTU format separated in z planes and time points (Fig. 6B) with partially lost corresponding metadata. Split datasets were reassembled and reshaped into 6D during import into napari using NFPP (Fig. 6C).

To date, sharing of FLIM data of various formats via OMERO is still challenged by the restricted availability of readers for specialized and less abundant formats and the limitation of dimensionality (Fig. 6F). Thus, while datasets in PTU format are not importable (Fig. 6C and F), those of LIF format lack visualization of detailed lifetime dimension but processed lifetime information and a Fast FLIM view are visualize (Fig. 6A and F). Datasets of Zarr and OME-TIFF formats are importable in 5D (Fig. 6C, D and F, Supplementary Fig. 2) with improved metadata annotation in OME-TIFF format. FLIM-based segmentation masks can enrich corresponding datasets in OMERO (Fig. 6E, Supplementary Fig. 2). The community-driven development of next-generation bioimage file formats (https://ome-model.readthedocs.io/en/stable/developers/6d-7d-and-8d-storage.html) will likely simplify the handling of FLIM datasets along the data life cycle including visualization, integrative lifetime analysis and sharing according to the FAIR principles in the future.

In summary, this study demonstrates the successful application of FLIM for lifetime unmixing of multiplexing fluorophores applied for live-cell imaging of dynamic processes in both *C. elegans* embryos and adult animals. Combining improved and transparent data handling and analysis workflows, this interdisciplinary project aims to broaden the application and the scientific readout of FLIM imaging in the life sciences.

## ACKNOWLEDGEMENTS

Imaging was performed at the Light Microscopy Facility at the Center for Molecular and Cellular Bioengineering (CMCB-LMF) of TU Dresden. We wish to acknowledge the support of the CMCB-LMF team, particularly Ellen Geibelt for technical support and maintenance of the microscope, and Silke Tulok (Core Facility Cellular Imaging at the Faculty of Medicine at TU Dresden) for support in bioimage data management using OMERO. We thank the Bio-Image Analysis Technology Development team of Physics of Life at TU Dresden (BiA-PoL) for provision of infrastructure and fruitful discussions, Tom Boissonnet (Heinrich Heine Universität Düsseldorf, I3D:bio) for support with metadata annotation and Josh Moore (German BioImaging - Gesellschaft für Mikroskopie und Bildanalyse e.V.) for feedback on the manuscript.

