## Supplementary figures 1 and 2 for "Fluorescence Lifetime Unmixing: A New Workflow for FLIM Live-Cell Imaging"

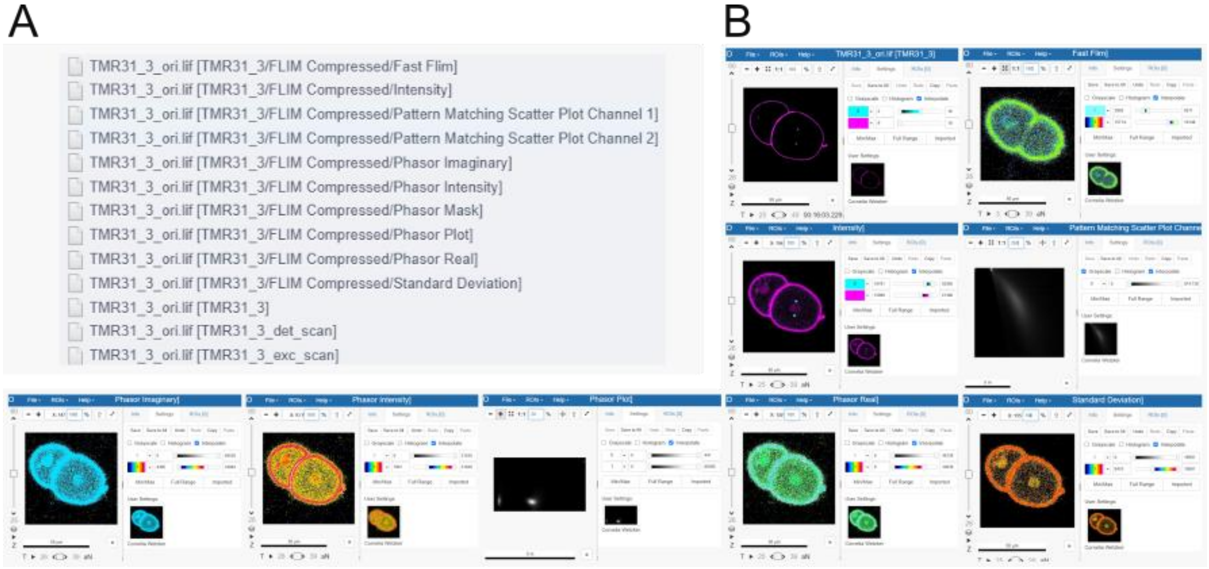


**Suppl. Fig. 1: Images of FLIM datasets in LIF format in OMERO.** (**A**) List of images as part of each imported 6D LIF dataset as visualized in OMERO. (**B**) Images show for example phasor parameters or FAST FLIM lifetimes, a single FLIM-based value per voxel, but no lifetime information as separate dimension. Look-up table settings are chosen for best visual contrast of the respective parameters for the datasets.

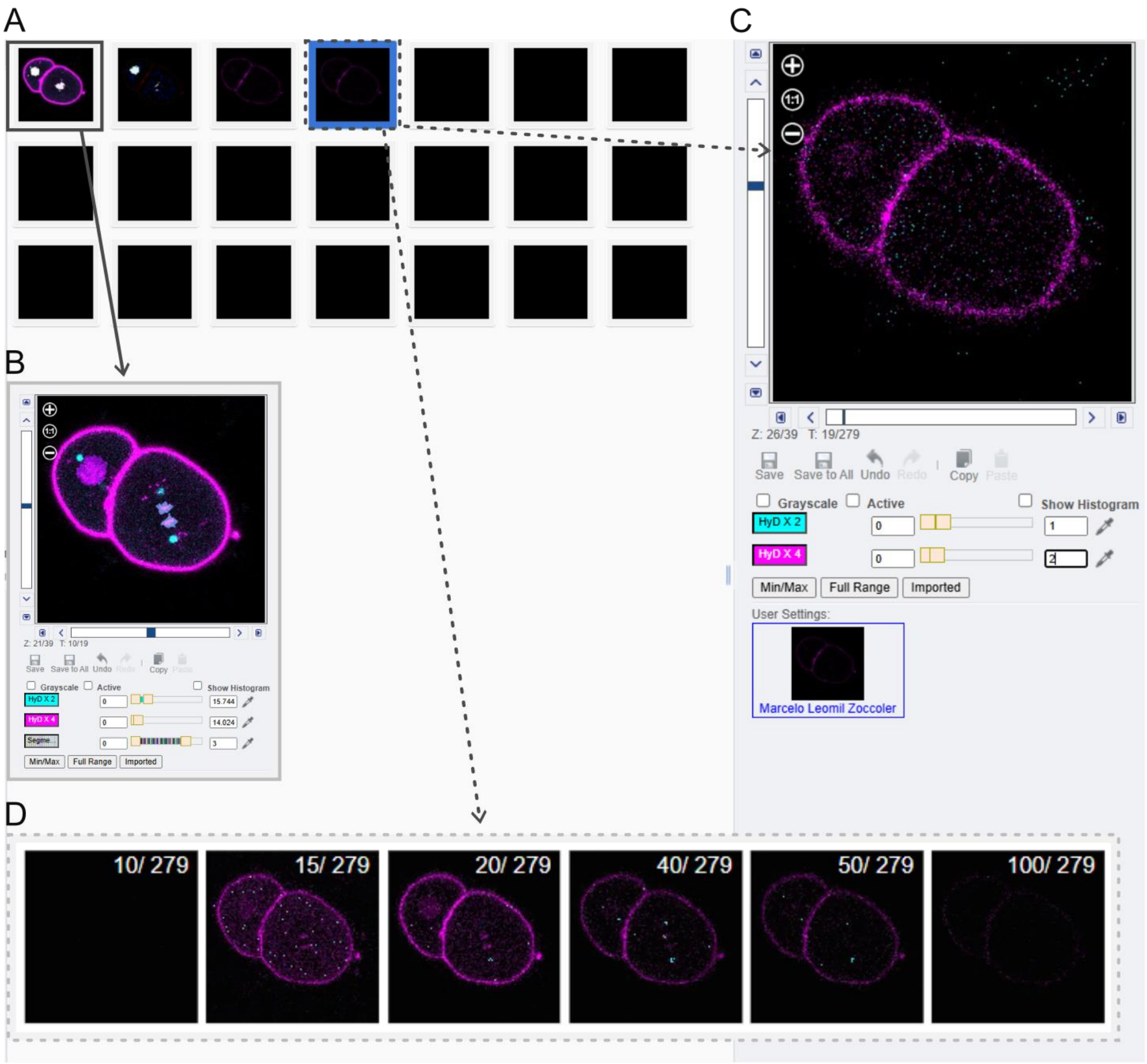


**Suppl. Fig. 2: Images of 5D FLIM datasets in OME-TIFF format in OMERO.** (**A**) Thumbnails of a 5D intensity image dataset with lifetime-based segmentation mask of chromatin as well as a 5D dataset with a lifetime slider, separated by time points (combined mCherry and mKate in magenta and centrosomes in cyan). (**B-D)** Detailed views of a single plane showing an exemplary 5D intensity dataset with NFPP-based segmentation of chromatin (**B**) and an exemplary single time point single plane of a 5D FLIM dataset (**C**) at different exemplary timepoints during exponential lifetime decay time points visualized in an OMERO.figure panel (**D**).
